# Atypical myxomatosis in European rabbits is caused by the recombinant myxoma virus involved in species jumping into hares

**DOI:** 10.64898/2026.07.25.740729

**Authors:** Jacqueline Carmona, Junior A. Enow, Ethan Ramsey, Sabeeha M. Reshi, Mackenzie Cashen, Ami D. Gutierrez-Jensen, Saige Munig, Natalie Reed, Kenneth Lowe, Jacquelyn Kilbourne, Grant McFadden, Simona Kraberger, Arvind Varsani, Masmudur M. Rahman

## Abstract

Myxoma virus (MYXV), a member of the *Leporipoxvirus* genus (species *Leporipoxvirus myxoma;* family *Poxviridae*), causes a highly lethal disease known as myxomatosis in European rabbits. In late 2018, a new natural MYXV isolate, MYXV-Tol (a.k.a. hare MYXV; ha-MYXV), emerged and caused myxomatosis-like disease with high mortality in Iberian hares, European brown hares, and European rabbits. This variant contains an approximately 2.8-kb insertion of a recombination cassette within the *M009L* gene encoding four additional genes, including the C7-like host range gene, *M159L*. M159 is essential for replication of MYXV-Tol in hare cells and is likely a key determinant of its pathogenicity in both hares and rabbits. Here, we compared the pathogenicity of wild-type MYXV-Tol (vMyx-Tol), an M159 deletion strain (vMyx-Tol-M159KO), and the classical MYXV-Lau strain (vMyx-Lau) in European rabbits. All three viruses caused systemic disease; however, vMyx-Tol and vMyx-Tol-M159KO produced clinical signs distinct from classical myxomatosis. Infection with vMyx-Tol and vMyx-Tol-M159KO was characterized by the absence of the typical primary and secondary nodular lesions, and caused severe edema, marked fluid accumulation, lymphocyte infection, and significantly reduced or no virus-neutralizing antibody responses. The disease caused by both vMyx-Tol and vMyx-Tol-M159KO progressed rapidly within 9-11 days, resulting in animals reaching humane euthanasia endpoints like vMyx-Lau. Deletion of M159 did not significantly alter MYXV-Tol pathogenicity in rabbits. Collectively, these findings demonstrate that MYXV-Tol has evolved to cause an atypical, amyxomatous-like acute to hyperacute disease in European rabbits and likely in hares.

**Significance:** Natural evolution enables viruses to cross species barriers and adapt to new hosts. Myxoma virus (MYXV), released in the 1950s in Australia and Europe as a biocontrol agent against European rabbits, became a classic model for real-time monitoring of virus evolution, virulence, and host adaptation. Although MYXV is typically host-restricted, a newly emerged natural isolate, MYXV-Tol, causes lethal disease in both hares and rabbits. Here, we show that MYXV-Tol induces an atypical, amyxomatous-like disease characterized by the absence of nodular lesions, severe edema, lymphocyte infection, and markedly reduced virus-neutralizing antibody responses. These findings reveal previously unrecognized virus–host interactions that shape disease outcome and provide new insight into the mechanisms driving viral adaptation and evolution.

## Introduction

Poxviruses (family *Poxviridae*) have a large double-stranded DNA genome and replicate in the cytoplasm of infected cells. Members of poxviruses infect humans, a wide variety of animals, and insects. Some poxviruses exhibit highly selected host-specific tropism and cause disease in that host, such as Variola, the cause of smallpox in humans; others have a broad host range and naturally infect multiple species, such as cowpox virus or mpox (1). Myxoma virus (MYXV; a member of the species *Leporipoxvirus myxoma*) is a highly host-restricted poxvirus that causes lethal disease called myxomatosis in European rabbits (*Oryctolagus cuniculus*). However, MYXV also causes asymptomatic infection in Brazilian cottontail rabbits (*Sylvilagus brasiliensis*) in South America and in California brush rabbit (*S. bachmani*) in the USA, where the virus has coexisted with these hosts for millennia (2–5). Myxomatosis disease in European rabbits was first described in 1898 in Uruguay when the virus, carried by mosquitoes, infected domestic European rabbits and caused fatal generalized disease (6). Subsequently, it was intentionally released in Australia and Europe in the 1950s as a biological agent for controlling the wild European rabbits, which became pests. Since its release on two different continents, the virus has undergone several genetic changes, leading to different virulence and disease outcomes (4, 7). Rabbits also co-evolved with MYXV and, on two different continents, independently developed resistance through identical sets of genetic changes (8–11).

Although rabbits and hares are members of the Leporidae family, hares were considered resistant to myxomatosis when MYXV was introduced in Europe in the 1950s. However, myxomatosis-like disease was only sporadically reported in the European brown hare (*Lepus europaeus*) caused by the type strain of MYXV, MYXV-Lau (12). A hare-adapted natural recombinant strain of MYXV, known as hare MYXV (ha-MYXV) or Toledo MYXV (MYXV-Tol), emerged in 2018 in Iberian hares (*Lepus granatensis*) on the Iberian Peninsula (13–15). This virus caused an initial epidemic in Iberian hares in several provinces of south and central Spain and later in Portugal (16, 17). During 2018-2020, the mean mortality rate was 55.4%, indicating high susceptibility of the Iberian hare to MYXV-Tol infection (16). The presence of the recombinant MYXV known as MYXV-Tol (or hare MYXV, ha-MYXV) was confirmed by diagnostic PCR; however, typical MYXV-Lau was not detected in hares. Although the genome of MYXV-Tol is highly similar to the parental MYXV-Lau strain released in Europe, it contains three disrupted genes, *M009L*, *M036L*, and *M152R*, and a ∼2.8-kb insertion on the left side of the genome, which is derived from an unknown poxvirus (13, 15). Later in 2020, the same virus had emerged in European brown hares, and since then, the virus has been detected in several European countries including Germany, the Netherlands, the Czech Republic, Slovakia, Hungary, and Italy (18–22). Sequencing of MYXV genomes from brown hares suggests that the virus isolates are closely related to MYXV-Tol, with few changes at the nucleotide level (20). MYXV-Tol also caused infection and death of wild and farm European rabbits throughout Europe and most recently in Algeria (20, 23, 24). Since mosquitoes are considered mechanical vectors of MYXV, the virus is expected to spread in hares and rabbits beyond Europe.

The MYXV-Tol recombinant cassette is inserted within the *M009L* gene, harboring four open reading frames (ORFs) phylogenetically related to MYXV genes encoding a poxvirus structural protein (*M157*), a thymidine kinase (*M158*), a C7-like (C7L) host range protein (*M159*), and a poly(A) polymerase subunit (*M160*), which are similar to the MYXV genes *M060R*, *M061R*, *M064R*, and *M065R*, respectively (13). We have recently reported that the C7-like host range protein M159 is essential for replication of MYXV-Tol in hare cell lines but not in rabbit cell lines (25). Deletion of M159 (vMyx-Tol-M159KO) also resulted in no replication in primary hare peripheral blood mononuclear cells (PBMCs), indicating that M159 is a key protein for MYXV-Tol replication in hare cells and is possibly linked with pathogenicity (25).

The deaths in hares and rabbits caused by MYXV-Tol are associated with severe swelling of the eyelids, face, ears, and genitals, accompanied by ocular purulent discharge. There is also development of distinct mucinous nodules and dermatitis across the face, limbs, and perineum. Based on the body conditions, it was anticipated that a shorter course of disease (10-14 days) would lead to mortality (13, 17, 23, 26). However, the disease progression and clinical outcomes have not been studied.

Here, we present the first report on the pathogenesis of MYXV-Tol in European rabbits in a laboratory setting. By comparing MYXV-Lau with MYXV-Tol and a host range mutant of MYXV-Tol, we can show that MYXV-Tol has naturally evolved to cause a distinct disease in European rabbits. Additionally, lacking the C7-like host range protein had no major change in the MYXV-Tol pathogenesis, suggesting that this new hare-adapted virus may have evolved in wild European rabbits before species jumping and causing lethal disease in hares by acquiring a novel recombination cassette containing the C7-like host range factor, M159.

## Results

### vMyx-Tol and vMyx-Tol-M159KO cause an atypical lethal disease in European rabbits

To evaluate the pathogenicity of wild-type MYXV-Tol (vMyx-Tol), Myx-Tol lacking the M159 gene (vMyx-Tol-M159KO), and the classical strain of wild-type MYXV-Lau (vMyx-Lau) in rabbits, New Zealand White rabbits were inoculated intradermally with these viruses. The clinical course of infection and disease progression was monitored daily until the rabbits reached the endpoint criteria and were humanely euthanized, which are summarized in Table 1. Rabbits infected with the vMyx-Lau strain developed typical nodular cutaneous myxomatosis characterized by the progressive development of a large, red, and raised primary skin lesion at the inoculation site by day 3, which became necrotic by day 4-5 (Table 1). However, rabbits infected with vMyx-Tol and vMyx-Tol-M159KO developed small, red, and mildly swollen primary skin lesions at the inoculation site by day 3-4, which eventually disappeared by day 6-7 and were no longer measurable (Table 1, Fig 1A and 1D). Rabbits infected with vMyx-Lau developed typical secondary lesions on ears, eyelids, nose, and other parts of the body by day 4-7 (Fig 1B). However, vMyx-Tol and vMyx-Tol-M159KO infected rabbits developed very small secondary lesions on ears by day 4-5 and eventually disappeared by day 6-7, as the swelling of ears increased (Fig 1B). Rabbits infected with vMyx-Lau had moderately swollen heads, eyelids, base of the ears, and anogenital areas after day 6-7 as the disease symptoms became severe. However, rabbits infected with vMyx-Tol and vMyx-Tol-M159KO had more severe overall swelling of heads, eyelids, ears, and anogenital areas after day 7 compared to vMyx-Lau (Fig 1E).

**Figure 1.**
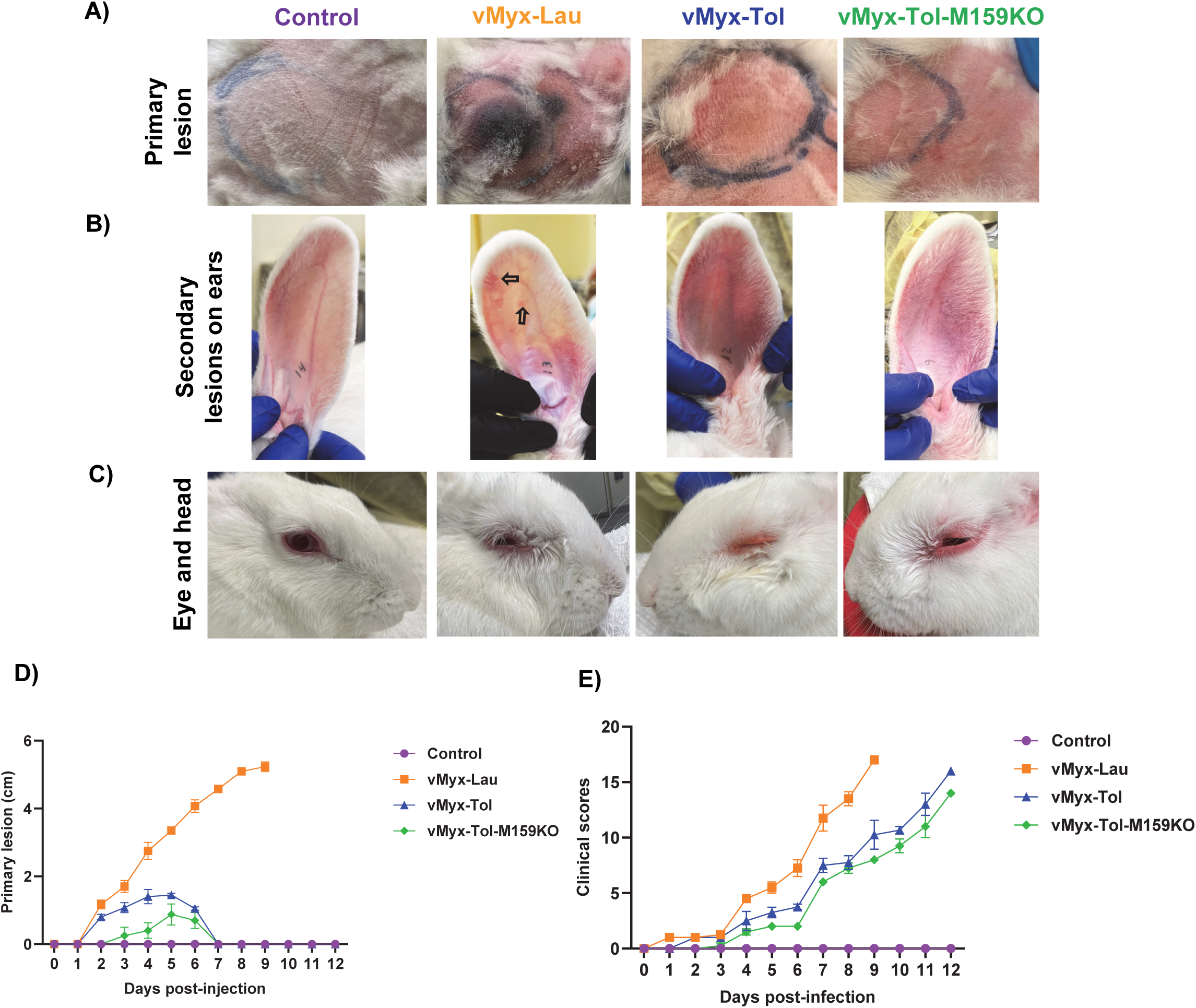
Clinical signs of disease in rabbits that are involved in the study. A) Pictures of primary lesion and injection area in the skin of the control and rabbits infected with the indicated three viruses on day 7 postinfection. B) Pictures of distal secondary lesions in the ears of the control and rabbits infected with the indicated three viruses on day 7 postinfection; arrow indicates secondary lesions in ears of vMyx-Lau-infected rabbits. C) Pictures of the eyes and head of the control and rabbits infected with the indicated three viruses on day 7 postinfection. D) The average size of the primary skin lesion of the rabbits injected with the indicated viruses. E) The average overall daily score for the rabbits injected with the indicated viruses.

**Table 1:** Pathogenesis of vMyx-Lau, vMyx-Tol, and vMyx-Tol-M159KO in New Zealand White rabbits

| **Day** | **vMyx-Lau** | **vMyx-Tol** | **vMyx-Tol-M159KO** |
| --- | --- | --- | --- |
| **0** | Four rabbits injected intradermally with $2 \times 10^3$ FFU | Four rabbits injected intradermally with $2 \times 10^3$ FFU | Four rabbits injected intradermally with $2 \times 10^3$ FFU |
| **1** | Mild redness at the injection site in all rabbits | No sign of reaction at the injection site | No sign of reaction at the injection site |
| **2** | Red and swollen primary lesion formation | Mild redness and swollen injection site in all rabbits | No sign of reaction at the injection site |
| **3** | Red and swollen primary lesion grew | Red and swollen small primary lesion | Mild redness and swollen injection site |
| **4** | Primary lesion red, swollen, and necrotic. Secondary lesions appear in ears and near primary lesion | Red and swollen primary lesion, secondary red spots near primary in two rabbits; | Red and swollen small primary lesion in all the rabbits |
| **5** | More secondary lesions appear in ears and near primary lesion, mild secretions from eyes, secondary lesions red and swollen; one rabbit has secondary lesion in eyes and nose | Primary lesion red and swollen; additional secondary red spots near primary in three rabbits; multiple red spots in ears for all the rabbits | Primary lesion red and mildly swollen, no secondary spots near primary lesion; multiple red spots in ears for all the rabbits |
| **6** | Primary lesions are very swollen and necrotic; Some of the secondary lesions are red, swollen, and necrotic; secondary lesions in eyes in more rabbits, mild secretions from eyes. One rabbit has profuse secretion from eyes. | Primary lesion remains the same; more secondary lesions like red spots in ears; ears are mildly swollen; mild secretions from eyes | Overall conditions remain unchanged; ears have more red spots. |
| **7** | Secondary lesions are red, swollen, necrotic; more secondary lesions in eyes and nose; ears, nose, head and genital area red and swollen for all rabbits; mild secretions from eyes; profuse secretions from eyes with reduced eating and drinking for one rabbit | Primary lesion disappears with swollen skin; in all the rabbit ears, eyes, nose, and genital area red and swollen; mild secretion from eyes; eating and drinking normal | Primary lesion disappears with swollen skin; in all the rabbit eyes, ears, nose, and genital area red and swollen; mild secretion from eyes; eating and drinking normal |
| **8** | Profuse secretions from eyes in all the rabbits; eyelids are closed; One of the rabbits was | Overall conditions remain the same; ears are very swollen from the base for some | Overall conditions remain the same; ears, eyes, nose, and genital area are more swollen |
|  | humanely euthanized due to disease severity | rabbits; attitude is quiet; reduced food consumption | for all rabbits; attitude is quiet; reduced food consumption |
| **9** | 3 rabbits were humanely euthanized due to severe disease symptoms | Condition of three rabbits remained the same with red and very swollen ears, eyes, nose, and genital area; mild to moderate secretions from eyes; one rabbit had profuse secretions from eyes, very swollen and closed; ears very swollen and stayed flat, nose, mouth, genital area very swollen; reduced drinking and eating and was euthanized. | Overall conditions remain the same; ears, eyes, nose, genital area more swollen with all rabbits; moderate secretion from eyes; attitude is quiet; reduced food consumption |
| **10** |  | Conditions remain stable with swollen overall body for three rabbits; reduced eating and drinking, quiet and less active | Overall conditions remain the same; very swollen ears, eyes, nose and genital area for all rabbits; attitude is quiet; reduced eating and drinking |
| **11** |  | Two rabbits had profuse secretions from eyes, heavily swollen and closed; ears heavily swollen from the base, nose, mouth and head heavily swollen; reduced drinking and eating; quiet and lethargic; two rabbits were humanely euthanized due to severe disease symptoms. | Two rabbit eyes were heavily swollen, closed with profuse secretions; ears, nose, mouth and genital area heavily swollen, reduced eating and drinking; quiet and lethargic; two rabbits were humanely euthanized due to severe disease symptoms. |
| **12** |  | The remaining two rabbits had profuse secretions from the eyes, heavily swollen and closed; ears heavily swollen from the base; nose, mouth, and head heavily swollen; reduced drinking and eating; quiet and lethargic; and the two rabbits were humanely euthanized due to severe disease symptoms. | The remaining two rabbits' eyes were heavily swollen, closed, and had profuse secretions; ears, nose, mouth, and genital area heavily swollen; reduced eating and drinking; quiet and lethargic; and the two rabbits were humanely euthanized due to severe disease symptoms. |

A clinical scoring system that has been reported in detail previously was used for daily evaluation of the progression of myxomatosis disease in these rabbits (27, 28). The average daily score from each group was plotted and used as a measure to reflect the disease progression (Fig 1E). A delayed development of clinical symptoms of disease was observed with the vMyx-Tol and vMyx-Tol-M159KO infected groups of rabbits compared to the vMyx-Lau infected rabbits. This difference was primarily due to the size of the primary skin lesion and absence of secondary lesions. However, as the disease progressed over time, other symptoms such as profuse secretion from the eyes, heavy swelling of the eyes, head, ears, and anogenital areas, decreased eating and drinking were observed after day 10 (Supp Fig S1). Eventually, all the rabbits infected with vMyx-Tol and vMyx-Tol-M159KO succumbed to severe disease and were humanely euthanized. Interestingly, the lack of C7-like protein, M159, had no effect on delaying disease progression compared to wild-type vMyx-Tol, supporting the prediction that M159 might have adapted to hare-specific host regulatory functions (25).

### vMyx-Tol causes viremia and edema in European rabbits

The normal rectal temperature for rabbits typically ranges between 38°C (100.4°F) and 40°C (104°F) with an average temperature of 38.9°C (102°F) (29). A rectal temperature below 37.9°C (100.2°F) is considered hypothermia, and a temperature of 39.89°C (103.8°F) or higher may indicate fever. In rabbits infected with vMyx-Lau, we regard rectal temperature >40°C (>104°F) as elevated and <40.22°C (<100.4°F) as indicating a poor prognosis (8). As shown in Fig. 2A, the temperature ranges in response to virus infection fluctuated with both vMyx-Lau and vMyx-Tol compared to the control rabbits. Although no hypothermia was observed with any of the rabbits, elevated temperature was observed with some of the rabbits infected with vMyx-Tol and vMyx-Tol-M159KO, indicating viremia (Fig. 2A).

**Figure 2.**
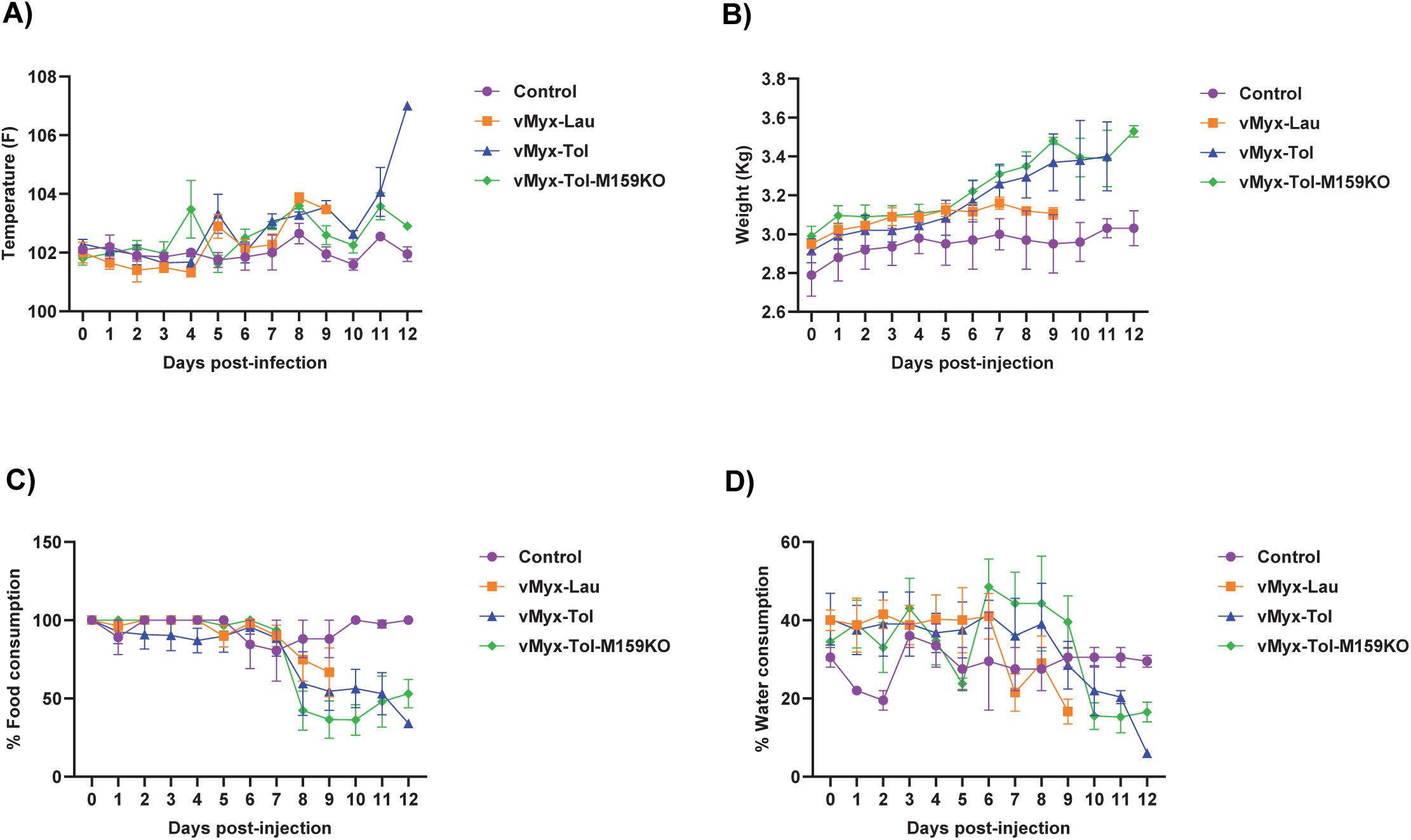
Rectal temperature, body weight, food and water consumption of rabbits infected with MYXV strains. The average daily rectal temperature (A), body weight (B), daily food intake (C), and daily water consumption (D) of control and rabbits infected with different viruses.

We measured the total body weight of the rabbits daily as one of the criteria for monitoring their health. We observed a consistent increase in weight for the control rabbits over time (Fig 2B). A similar trend was observed with the virus-infected rabbits until day 6. After this time, vMyx-Lau-infected rabbits didn’t gain weight until euthanized due to reduced eating and drinking from the disease symptoms. Surprisingly, rabbits infected with both vMyx-Tol and vMyx-Tol-M159KO viruses consistently gained weight until euthanized, although these virus-infected rabbits started showing severe disease symptoms (Fig 1E and Fig S1). Comparing day 0 and 9, we observed that vMyx-Tol and vMyx-Tol-M159KO-infected rabbits gained 15-16% in weight, whereas control and vMyx-Lau-infected rabbits gained only 5-6% in weight (Fig 2B). Additionally, we noticed a 10-15% increase in water consumption between days 6-9 for vMyx-Tol and vMyx-Tol-M159KO-infected rabbits, which might have allowed fluid retention in these rabbits. However, starting on day 8, food consumption decreased by 10-25% after day 8 with vMyx-Tol- and vMyx-Tol-M159KO-infected rabbits as the disease severity increased (Fig 2C and 2D). Thus, the likely cause of weight gain is due to the accumulation of fluid in the body.

Hematoxylin and eosin-stained sections of skin from the inoculation site were examined after euthanasia. Animals inoculated with all three viruses showed widespread swelling and vesicular degeneration of the epidermis, and massive disruption of the dermis compared to the control rabbit skin tissues (Fig S2). There was more infiltration of myxomatous stroma in the dermis of vMyx-Lau infection compared to the vMyx-Tol infection (Fig S2A). Although there was no typical primary lesion at the injection site of vMyx-Tol and vMyx-Tol-M159KO, there was substantial disruption of the epidermis and dermis compared to the control rabbit skin (Fig S2B-D).

### vMyx-Tol and vMyx-Tol-M159KO-infected rabbit serum had lower levels of virus neutralizing antibodies compared to vMyx-Lau

Since vMyx-Tol and vMyx-Tol-M159KO-infected rabbits showed different clinical symptoms and disease progression than vMyx-Lau, we collected whole blood during euthanasia and prepared serum to measure the level of virus neutralizing antibodies (Nabs) and circulating cytokines using a plaque reduction neutralization assay (PRNT) and cytokine array, respectively. Our PRNT assay results demonstrate that serum from vMyx-Lau-infected rabbits neutralized all three tested viruses between 60-100% at the lowest 10-fold dilution. However, after 10^3^-fold dilution, the level of neutralization significantly dropped (Fig 3A). A similar assay with antibodies from a mutant MYXV (vMyx-M130KO virus) which did not cause lethal disease in rabbits, resulted in a relatively higher titer of neutralizing antibodies (data not shown). Unlike vMyx-Lau-infected rabbit serum, serum from control, vMyx-Tol, and vMyx-Tol-M159KO-infected rabbits showed very low or no neutralization against any of the three viruses even at 10-fold dilution (Fig 3B, 3C, and 3D). To further test the reactivity of the antibodies against viral proteins, we performed Western blot analysis using purified virions. The results indicate that antibodies against vMyx-Lau-infected rabbits detected similar virion proteins present in vMyx-Lau and vMyx-Tol virions. However, antibodies against vMyx-Tol and vMyx-Tol-M159KO-infected rabbits detected very few virion proteins with different sizes compared to vMyx-Lau (Supp Fig S2).

**Figure 3.**
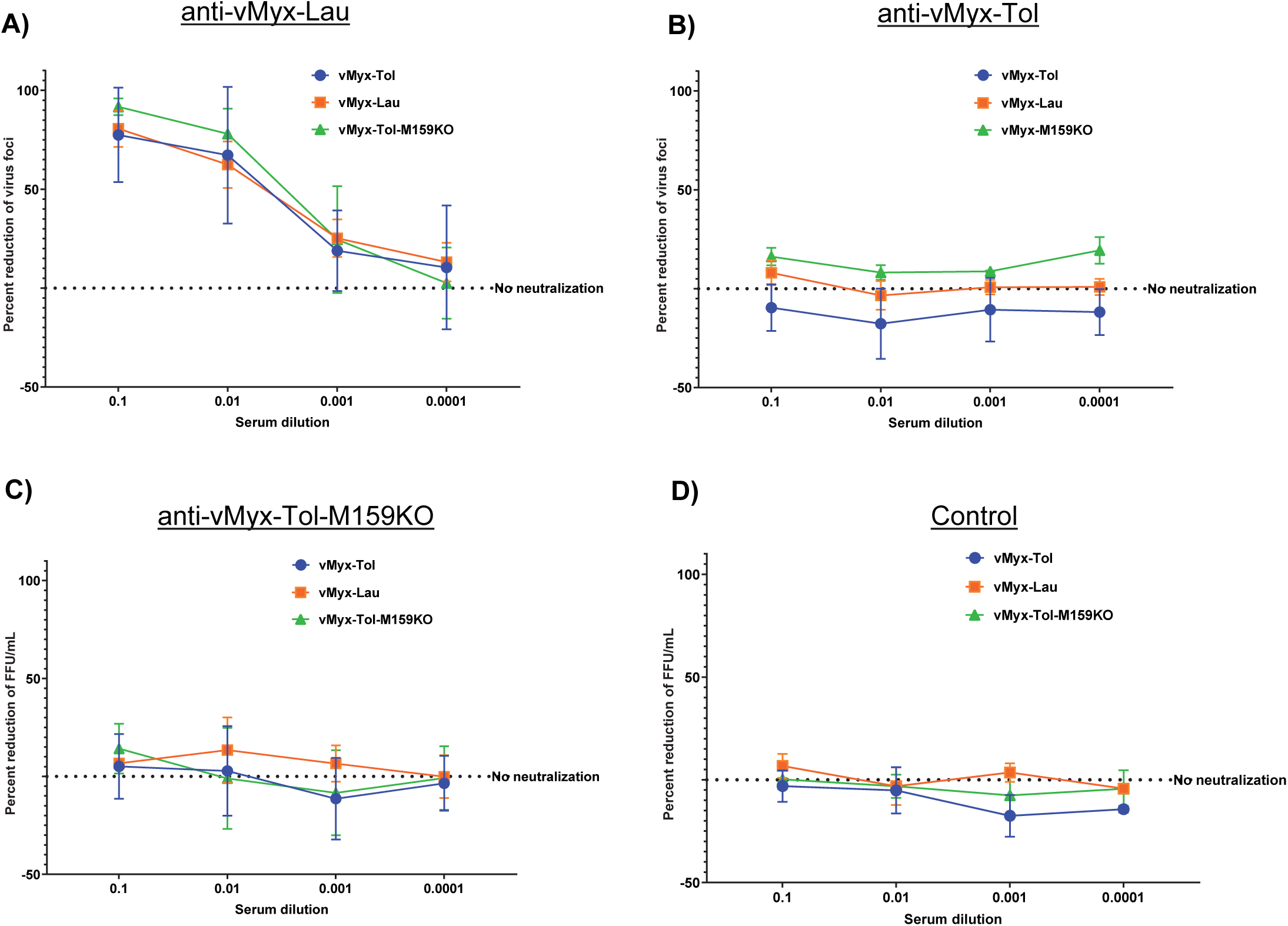
Evaluation of virus neutralizing ability of the serum collected from the rabbits infected with different viruses. PRNT assay with sera from rabbits infected with A) vMyx-Lau, B) vMyx-Tol, C) vMyx-Tol-M159KO, and D) uninfected rabbits against the indicated viruses is shown.

Furthermore, we isolated PBMCs from the blood that were collected before euthanasia of rabbits. The cells were observed under a fluorescence microscope, and it was seen that the majority of PBMCs from rabbits infected with vMyx-Tol and vMyx-Tol-M159KO were infected with virus, based on the expression of reporter GFP and TdTomato protein, whereas PBMCs from rabbits infected with vMyx-Lau had no or very few virus-infected cells (Fig S2D).

## Elevated expression of selected cytokines in vMyx-Tol infection

To test whether there are differences in immune responses against these different virus infections, we analyzed the level of selected inflammatory cytokines in the serum collected at the end of the study using a commercial rabbit cytokine array (RayBiotech). The cytokines are IL-1α, IL-1β, IL-8 (CXCL8), IL-17A, IL-21, Leptin, MIP-1β (CCL4), MMP-9, NCAM-1 (CD56), and TNFα (Fig 4). We observed an increased level of IL-17A, IL-1α, and Leptin in rabbits infected with vMyx-Tol compared to the control (Fig 4A, 4D), whereas the NCAM-1 level was significantly higher in rabbits infected with vMyx-Tol-M159KO (Fig 4B). Increased level of cytokines IL-8, MMP-9, and MIP-1β was seen with vMyx-Lau infection compared to the control rabbits (Fig 4A, 4C). Elevated levels of these cytokines indicate chronic inflammation with these viruses under different infection conditions and recruitment of immune cells.

**Figure 4.**
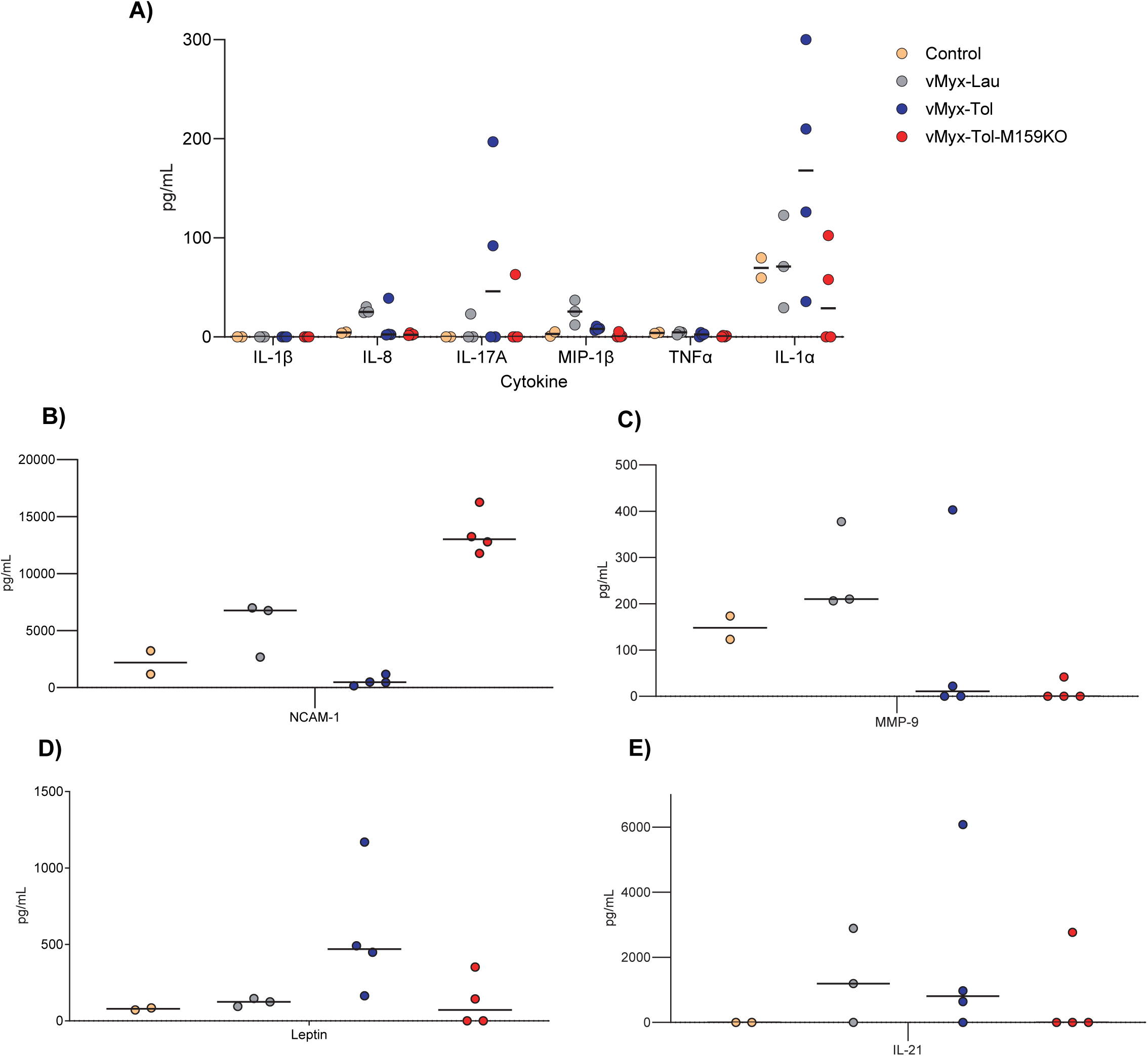
Infection with different MYXV strains induces changes in the level of circulatory cytokines. Serum was prepared from blood collected during euthanasia. Using a commercial rabbit cytokine array, the level of 10 different cytokines was measured.

Using RT-qPCR, we measured the changes in the expression level of selected cytokines in the primary skin lesion of rabbits infected with these viruses. Compared to vMyx-Lau infection, the level of proinflammatory cytokines such as TNFα, IL-1β, and IFNɣ was higher with vMyx-Tol and vMyx-Tol-M159KO virus infection (Fig S4). However, IL17 was higher with vMyx-Lau infection. Elevated level of these cytokines again indicates relatively higher inflammation with virus infection compared to the control rabbits.

## Discussion

Poxviruses undergo rapid evolution for host adaptation, using diverse mechanisms such as gene duplications, gene loss, homologous and nonhomologous recombination, and horizontal gene transfer (30, 31). These evolutionary mechanisms drive poxvirus host species jumps and adaptation to a new host, such as in the case of mpox, cowpox virus, and MYXV (32, 33). Following the release of MYXV in Australia and Europe in the 1950s, the virus has undergone rapid genetic changes, creating divergent lineages to bypass rabbit immunity, which resulted in a major shift in virulence within a few years after the release. Based on those genetic changes, the virus isolates were graded depending on the level of virulence in rabbits (34). Interestingly, these genetic changes in the virus happened across both continents, resulting in parallel, convergent evolution where disease phenotype is matched, but the exact genotypes differ (7, 35, 36). Even with those genetic changes, until 2018, MYXV was known to be restricted to rabbits. However, the new MYXV-Tol isolate acquired a recombination cassette with additional genes, allowing the virus to infect both hares and rabbits. The recombination cassette contains four genes, including a C7-like host range protein-encoding ORF *M159L*. We reported that deletion of *M159L* from MYXV-Tol made the virus defective in replication in hare cell line and primary hare PBMCs (25). The classical wild-type MYXV-Lau was also defective in replication in hare cells, which can be reversed by the expression of M159 protein from MYXV-Lau (25). Additionally, M159 was not required for MYXV-Tol replication in rabbit cells. These results indicated that the C7-like protein in the recombination cassette probably allowed the virus to adapt to hares and cause disease in both hares and rabbits.

Since MYXV-Tol causes lethality in wild and farm rabbits, it is important to understand the disease in greater detail. So far, based on the necropsy of dead hares, the pathogenesis of MYXV-Tol results in acute or hyperacute myxomatosis, characterized by generalized edema, congestion, and hemorrhages (14). In this study, we inoculated New Zealand White rabbits intradermally with a virus dose that causes lethality by MYXV-Lau, and the disease progression and clinical symptoms have been documented (27, 37). At the inoculation site, typically MYXV-Lau causes a necrotic primary lesion within 4-5 days; surprisingly, MYXV-Tol and MYXV-Tol-M159KO formed a small red and swollen primary lesion, which disappeared over time by day 7. The typical secondary lesions in ears, eyes, nose, and near the primary lesion caused by MYXV-Lau did not develop with MYXV-Tol or MYXV-Tol-M159KO. Only small red spots that formed as an indication of secondary lesions quickly disappeared due to the edema and as the disease progressed. The absence of myxomatosis-associated tumor-like lesions in rabbits and hares was reported in a study where they evaluated the efficacy of anti-MYXV vaccines against MYXV-Tol or in wild natural infection with MYXV-Tol (26). Myxomatosis lesions were not always detected in wild hares and rabbits that were found dead with MYXV-Tol infection. Whether the absence of nodular primary and secondary lesions is linked to genetic changes needs to be further investigated. Our results show that the clinical symptoms associated with myxomatosis, such as swelling of the nose, face, base of the ears, anogenital area, mild to heavy discharge from the eyes and nose, were delayed by only two days with MYXV-Tol and MYXV-Tol-M159KO. However, once the symptoms started, the disease progressed rapidly and required humane euthanasia of the animals. The most notable difference compared to MYXV-Lau was the edema with both MYXV-Tol and MYXV-Tol-M159KO, resulting in swelling of the different body parts, which also increased the body weight. Based on these clinical symptoms and rapid disease progression (9-12 days after virus injection), MYXV-Tol would correspond to a Grade I virus according to the classic MYXV virulence grade (34, 36). Clearly, the absence of *M159L* in MYXV-Tol had very little impact on the pathogenicity of MYXV-Tol in European rabbits.

Our findings that MYXV-Tol had only a minimal effect on disease progression compared to the parental vMyx-Lau strain were unexpected, since vMyx-Tol has several gene disruptions. For example, the insertion of the recombination cassette in MYXV-Tol resulted in the disruption of the *M009L* gene, the role of which in MYXV pathogenicity is still unknown. Apart from *M009L*, two other genes, *M036L* and *M152R*, are also disrupted/truncated in MYXV-Tol (13). Our results indicate that disruption of these genes might have no or very little effect on MYXV-Tol pathogenicity. Mutations in these three genes are also reported in the MYXV isolates from Australia and Europe that were collected and sequenced after the release of parental MYXVs. *M009L* ORF, a ubiquitin ligase, is disrupted in almost all isolates from Australia sampled after 1990, in some European isolates, and in the California MSW strain of MYXV (35, 38). However, these mutations were predicted to have no effect on virulence, since they were also present in isolates with the highest virulence (35). Presence of mutations in the *M036L* ORF has been reported in several field isolates from the UK and Australia showing different levels of virulence (7, 35, 38). Interestingly, MYXV isolates that have *M009L* and *M036L* mutations demonstrated amyxomatous (non-nodular) disease with different levels of virulence (35). *M152R* ORF encodes a serine proteinase inhibitor (Serp3), which has been shown to be associated with MYXV virulence (39). In MYXV-Tol, this gene is disrupted by insertion of an early stop codon (13). *M152R* ORF is also disrupted in the California MSW strain of MYXV (40). In the future, the role of M152 associated with changes in MYXV-Tol pathogenicity needs to be further explored.

Before the emergence of MYXV-Tol in hares, antibodies against MYXV have been reported in various hare populations in Europe, indicating exposure to MYXV-Lau, although they were resistant to the virus (41). However, outbreaks of MYXV-Tol resulted in significantly enhanced seroprevalence (more than 24.7%) (41). There are mixed reports on detection of anti-MYXV antibodies in hares. For example, in a road-killed European hare which was MYXV-positive, anti-MYXV antibodies were not detected (21). Similarly, very low antibody titers were detected in sera from wild rabbits with infection against the MSW strain of MYXV (42). These reports led us to test the level of neutralizing antibody in the rabbits from the serum collected at the terminal disease stage before euthanasia. We observed significantly lower virus neutralizing antibody titers in rabbits infected with MYXV-Tol and MYXV-Tol-M159KO compared to vMyx-Lau. Our results thus support the findings from the infections in the field that MYXV-Tol results in less anti-MYXV antibody. But this observation needs to be studied in greater detail to understand this phenomenon with vMyx-Tol infection. Furthermore, we observed that PBMCs collected from the blood of terminal infection were highly infected in the case of MYXV-Tol and MYXV-Tol-M159KO viruses, indicating a possible different route of virus spread compared to MYXV-Lau, which may result in the infection of immune cells that are involved in antibody production. Another possibility is that vMyx-Tol infects CD4^+^ T cells, impairing its ability to present antigen and promote B cell isotype switching and maturation, which generates a low-affinity polyclonal population that is ineffective at neutralizing the virus. Viruses such as human immunodeficiency virus, cytomegalovirus, and Epstein-Barr virus (43).

MYXV can spread in different organs to cause systemic diseases (42, 44). In previous studies and naturally found dead hares and rabbits, the viral loads of MYXV-Tol were reported to have similar levels in different organs (45). High viral load was detected in the lungs of rabbits compared to hares; this could be due to the acute infection in rabbits (26, 46). To compare these two viruses, future studies need to focus on tracking how the virus is progressing from the inoculation site to different organs and causing a systemic disease.

Overall, based on our results the disease phenotypes caused by MYXV-Lau and MYXV-Tol can be divided into two distinct categories: (i) a nodular cutaneous or “myxomatous” disease with primary lesion at the inoculation site and secondary cutaneous lesions on different parts of the body caused by classical MYXV-Lau, and (ii) a disease that resembled the “amyxomatous” phenotype characterized by a poorly defined primary lesion and very few poorly transient secondary lesions by MYXV-Tol. MYXV-Tol infection was also accompanied by very swollen heads, ears, eyelids, and perineum compared to MYXV-Lau. Reduced antibody titer and spread of virus were different. These results indicate that vMyx-Tol may have adapted in rabbits to cause a different type of disease before jumping to hares.

## Materials and Methods

### Cell lines and viruses

The different recombinant viruses used in this study are: vMyx-Tol (vMyx-Tol-TdT-GFP), a wild-type MYXV-Tol expressing reporter tdTomato (TdT) fluorescent protein under the control of a poxvirus late p11 promoter and green fluorescent protein (GFP) under the control of a poxvirus synthetic early-late promoter (25). vMyx-Tol-M159KO (vMyxTol-ΔM159-TdT-GFP), a recombinant MYXV-Tol lacking the *M159L* gene (25). vMyx-GFP, a wild-type MYXV-Lau expressing GFP under a poxvirus synthetic early-late promoter (37). All these viruses were amplified using the rabbit RK13 cell line grown in Dulbecco’s Modified Eagle Medium (DMEM), supplemented with 10% fetal bovine serum and 1% penicillin/streptomycin.

### Animal study

New Zealand White rabbits were purchased from Charles River Laboratories International, Inc (USA). The animal study was approved by the Institutional Animal Care and Use Committee (IACUC) at Arizona State University, and studies were performed as described before (27, 37). 2000 focus forming units (FFU) of the tested virus were resuspended in 100 µL of phosphate-buffered saline (PBS) and inoculated intradermally in the right flank of each rabbit after shaving the area with a clipper. Daily physical examinations were conducted to evaluate the rabbits’ health and condition by monitoring weight, rectal temperature, food and water intake, attitude, hydration status, urine and feces output, posture, and indications of primary lesions and appearance of secondary lesions. Based on these evaluations, the rabbits received a daily clinical score. The animals were humanely euthanized when the clinical score reached endpoint criteria, or animals had open-mouth breathing due to respiratory stress, orthopnea, cyanosis, or no food and water intake for 48 h.

### Plaque reduction neutralization test (PRNT) with rabbit serum

To evaluate the neutralization capacity of antibodies generated after inoculation with vMyx-Lau, vMyx-Tol, and vMyx-Tol-M159KO, plaque reduction neutralization test (PRNT) assays were performed using serum collected after euthanizing the rabbits. Serum was separated from blood using serum separator tubes (BD Microtainer™) after high-speed centrifugation (13,000 g, 10 min at 4°C). The serum was inactivated for 30 minutes at 56°C and stored at -80°C until needed. Four serial dilutions (1:10, 1:10^2^, 1:10^3^, 1:10^4^) of serum (45 µL) were incubated with 10 µL (100 FFU) of virus for 1 hour at room temperature (flicking the tubes every 15 minutes). The final volume of virus-serum mix was brought up to 500 µL and added to a monolayer of RK13 cells that were plated in a 12-well plate 24 hours prior (80-90% confluency, about 4 × 10^5^ cells per well). After 1 hour at 37°C, the serum-virus mixture was aspirated, and cells were overlaid with agarose-containing medium (1% low-melting agarose in DMEM with 10% FBS) and incubated for 48hours at 37°C. Viral fluorescence foci were counted using a fluorescent microscope, and data were represented as the percent reduction of foci compared to the virus-only control by the reciprocal of serum dilution. Every serum dilution was tested in triplicate and compared to neutralization with serum from PBS-inoculated rabbits.

### RT-qPCR

To measure different cytokine transcript levels, tissue samples from the primary lesion were collected into RNAlater solution (Invitrogen) and stored at -80°C until used. Tissue samples were dissected with scissors or a scalpel into smaller fragments of 30-60 mg and homogenized (VWR Mini Bead Mill) with ceramic beads (1.4 mm) in 1 mL of SNAP-60 (Amsbio) reagent. RNA was isolated according to the manufacturer’s instructions. The total RNA was reverse transcribed to complementary DNA (cDNA) using the High-Capacity cDNA Reverse Transcription Kit (Applied Biosystems) and stored at -20°C until needed. The equivalent of 100-200 ng of RNA was used per qPCR reaction in a total volume of 10 µL that included primers (0.2uM each) and 5 µL of SYBR Green mix (Applied Biosystems). Reactions were run in triplicate on a QuantStudio 3 (Applied Biosystems) (2 min 60°C, 10 min 95°C, 40 cycles of 15 sec at 95°C, 60 sec at 60°C). A dissociation curve (15 sec at 95°C, 15 sec at 60°C, and 15 sec at 95°C) was run for quality control, and the results were analyzed with Microsoft Excel. Gene expression was quantified with the comparative Ct method, where the gene expression of each sample was normalized to the expression of the housekeeping glyceraldehyde-3-phosphate dehydrogenase (GAPDH) gene and then compared to the normalized gene expression of the uninfected control rabbit. The relative fold change in gene expression was plotted as the Log_2_(Δ-ΔC_t_) of the difference between the infected and the uninfected control using GraphPad Prism software. Primer sequences used in this experiment are shown in Supplemental Table S1.

### Histology

Healthy skin from uninfected rabbits and primary lesions formed on the skin at the injection site were dissected from rabbits infected with vMyx-Lau, vMyx-Tol, and vMyx-Tol-M159KO. Samples were submerged in 10% neutral buffered formalin for at least 48 hours. Tissue samples were transferred into 70% ethanol solution and stored at 4°C in the dark until needed and processed within 2 weeks. Tissues were trimmed and fitted into sample cassettes and dehydrated with graded ethanol baths (70% to 100%) and cleared with Xylene using the HistoCore PEARL Automated tissue processor (Leica Biosystems). The tissue was paraffin-embedded with the HistoCore Arcadia Embedding center (Leica Biosystems) and cut on a HistoCore BIOCUT R Mechanical rotary microtome (Leica Biosystems) into 5 µm sections that were placed on a warm water bath and picked up onto Hareta charged slides (Springside Scientific) to dry overnight at room temperature. Slides were stained with Hematoxylin and Eosin (H&E) using an ST5010 Autostainer XL automated slide stainer (Leica Biosystems) at the Regenerative Medicine Core facility (Arizona State University). Slides were imaged and processed using the Olympus VS200 slide scanner, ImageJ, and OlyVIA software v5.1.

### Multiplex cytokine array

Cytokine quantification was performed using the Quantibody Rabbit Cytokine Array 1 (RayBiotech) according to the manufacturer’s instructions. The microarray consists of a slide divided into 16 wells, and each well has an array of capture antibodies bound to the glass surface. Each antibody is printed in quadruplicate, and each slide has 8 wells dedicated to the 7-dilution cytokine standard mix provided and a cytokine-free negative control. After blocking, serum samples from PBS- and MYXV-infected rabbits were diluted by 2-fold and applied to the assigned well and incubated with gentle rocking overnight at 4°C. Samples were decanted, and the wells were washed 5 times at room temperature and gentle rocking using the buffer I provided and then 2 more times with buffer II. Wells were probed with the biotinylated secondary antibody for 2 hours at room temperature and washed 5 times with buffer I and 2 times with buffer II. Wells were incubated with a Cy3 equivalent dye-conjugated streptavidin for 1 hour at room temperature and washed again. Slides were placed in a slide washer/dryer tube and washed for 15 minutes with buffer I and 5 minutes with buffer II with gentle rocking. Slides were carefully dried by gentle centrifugation (1000 rpm, no acceleration, no brakes) and removal of remaining droplets with a pipette tip. Samples were imaged with a laser scanner using the green channel (Cy3), and the data were analyzed using the company’s Q-Analyzer software (RayBiotech) and visualized with GraphPad Prism.

### Western blot

Equal amounts of total protein (quantified by BCA assay-Fisher Scientific) from viral stocks of WT-MYXV (MYXV Lausanne), MYXV strain Toledo (MYXV-Tol), and a MYXV-Tol-M159ko were loaded onto 4-20 % Miniprotean TGX gels (BioRad) and run at 110V until the protein standard ladder was resolved. Proteins were transferred onto a polyvinylidene difluoride (PVFD) membrane using the Power Blotter station (Invitrogen). PVDF membranes were blocked with 5% nonfat milk in Tris-Buffered Saline with Tween 20 (TBST) buffer (20mM Tris, 150 mM NaCl, 0.1% Tween 20, pH 7.6) for 1 hour at room temperature. Proteins were probed with serum (1:100) from infected (vMyx-Lau, vMyx-Tol, vMyx-Tol-M159KO) or uninfected rabbits at 4°C overnight. Membranes were washed three times (10 minutes each) with ice-cold TBST and incubated with Goat anti-Rabbit IgG antibody conjugated to horseradish peroxidase (HRP) in TBST with 5% nonfat dry milk for 1 hour at room temperature. Membranes were washed three times again with ice-cold TBST and developed using the Immubilon Western substrate (Millipore); chemiluminescent signal was detected with an ImageQuant 800 imager (Amersham).

### Ethics statement

This animal study was performed following the IACUC protocol number 23-1969R and titled “Studies in poxvirus host range genes and tropism”. IACUC protocol was approved by the Arizona State University Animal Care Services in accordance with the guidelines set by the Association for Assessment and Accreditation of Laboratory Animal Care (AAALAC).

## Statistical analysis

Statistical analyses were performed using GraphPad Prism 10.5.0 software (La Jolla, CA). Values are represented as the mean ± SD for at least two or three independent experiments. The two-tailed unpaired Student’s *t*-test was used to compare two independent groups, and the one-way ANOVA was used to compare three or more groups. *P* values are reported as follows: not significant (ns) *P* > 0.05, * *P* < 0.05, ** *P* < 0.01, *** *P* < 0.001, **** *P* < 0.0001.

## Acknowledgements

We acknowledge Arizona State University Department of Animal Care and Technologies (DACT) staff members for their support with animal research.

## Data availability

The data that support the findings of this study are available in the article.

---

**Figure S1.**
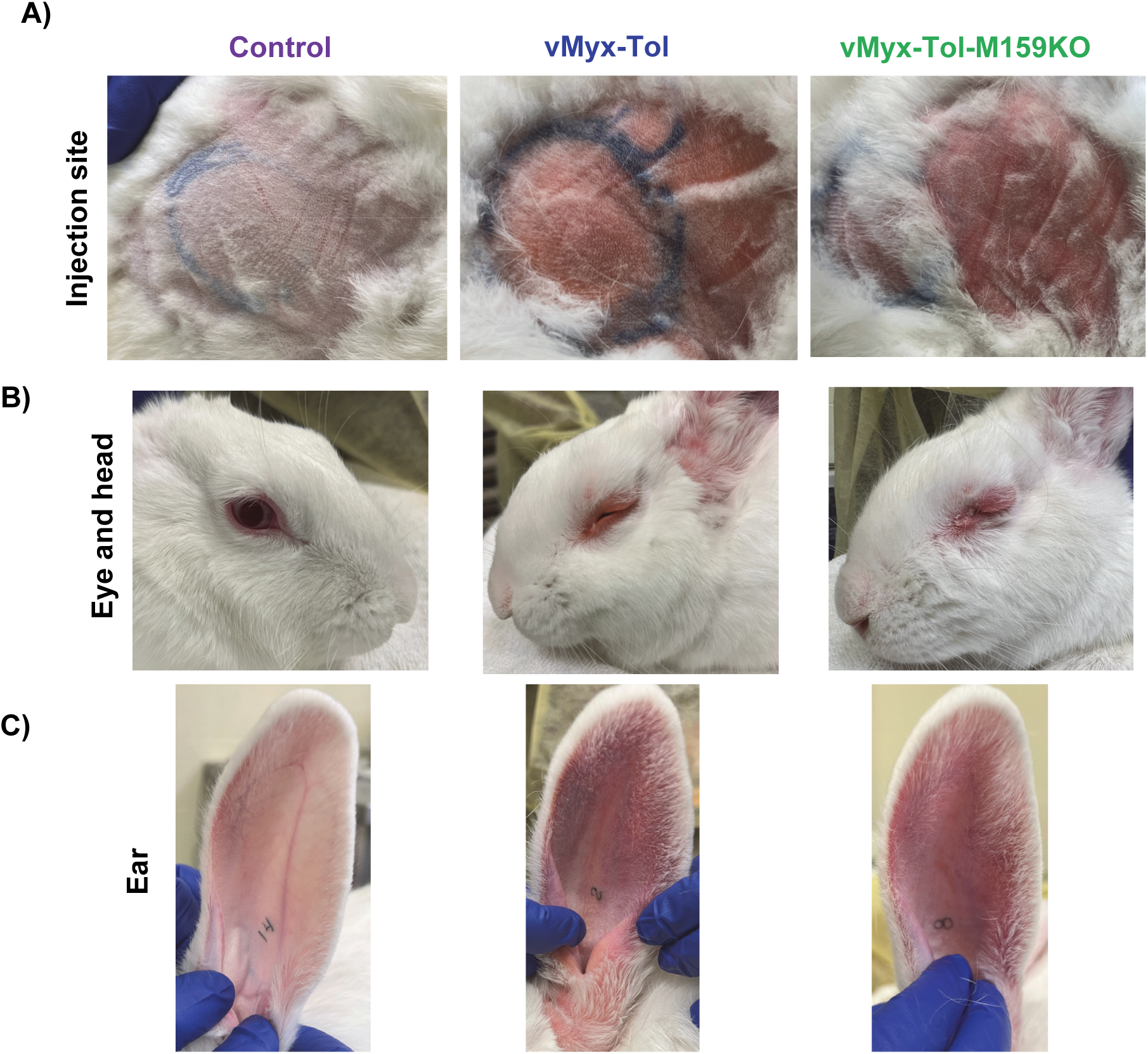
Clinical signs of disease in rabbits on day 11. A) Pictures of the injection area in the skin of the control and rabbits infected with the indicated viruses on day 11 postinfection. B) Pictures of the eyes and head of the control and rabbits infected with the indicated viruses on day 11 postinfection. C) Pictures of severely swollen ears of the rabbits infected with the indicated viruses on day 11 postinfection.

**Figure S2.**
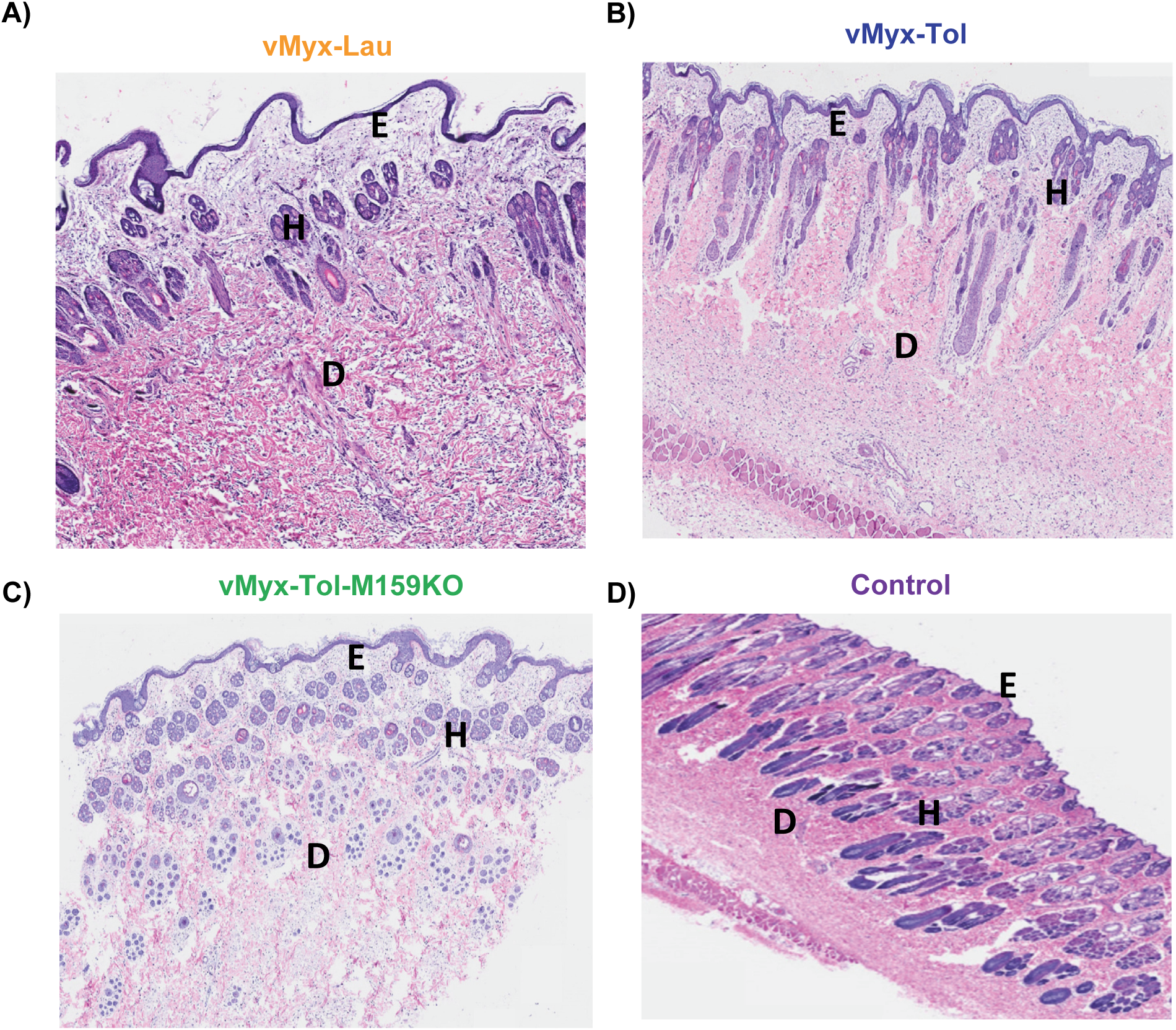
Histopathology analysis of rabbit skins collected from the injection site. Representative primary skin lesion from rabbits infected with A) vMyx-Lau, B) vMyx-Tol, C) vMyx-Tol-M159KO, and D) control rabbit. E, epidermis; D, dermis; H, hair follicle.

**Figure S3.**
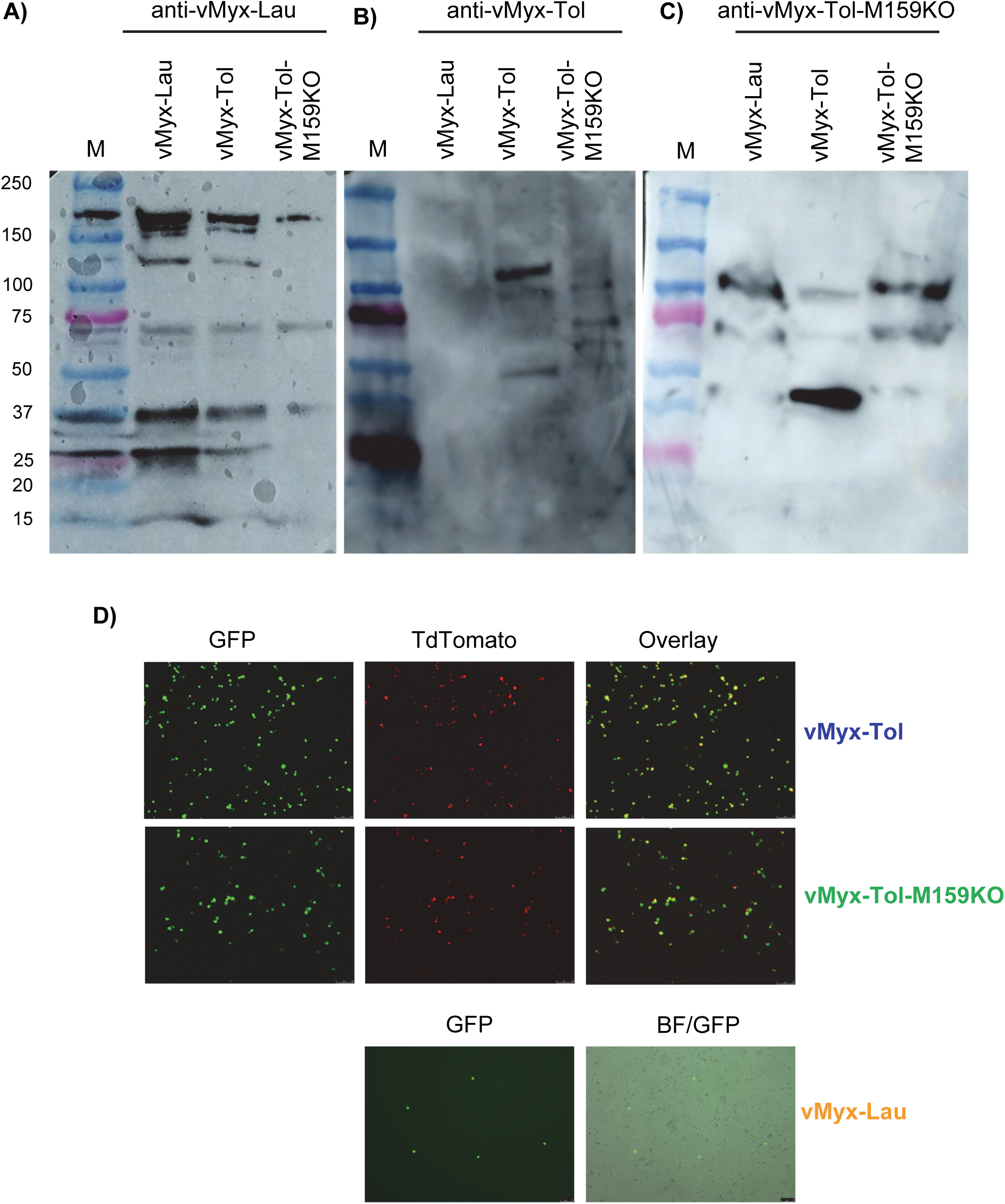
Detection of viral proteins using serum from the rabbits infected with the indicated viruses. Western blot analysis to detect the proteins from the indicated purified viruses using the serum from A) vMyx-Lau, B) vMyx-Tol, C) vMyx-Tol-M159KO, and D) Fluorescence microscopic images of PBMCs that were isolated from the blood of the indicated virus-infected rabbits.

**Figure S4.**
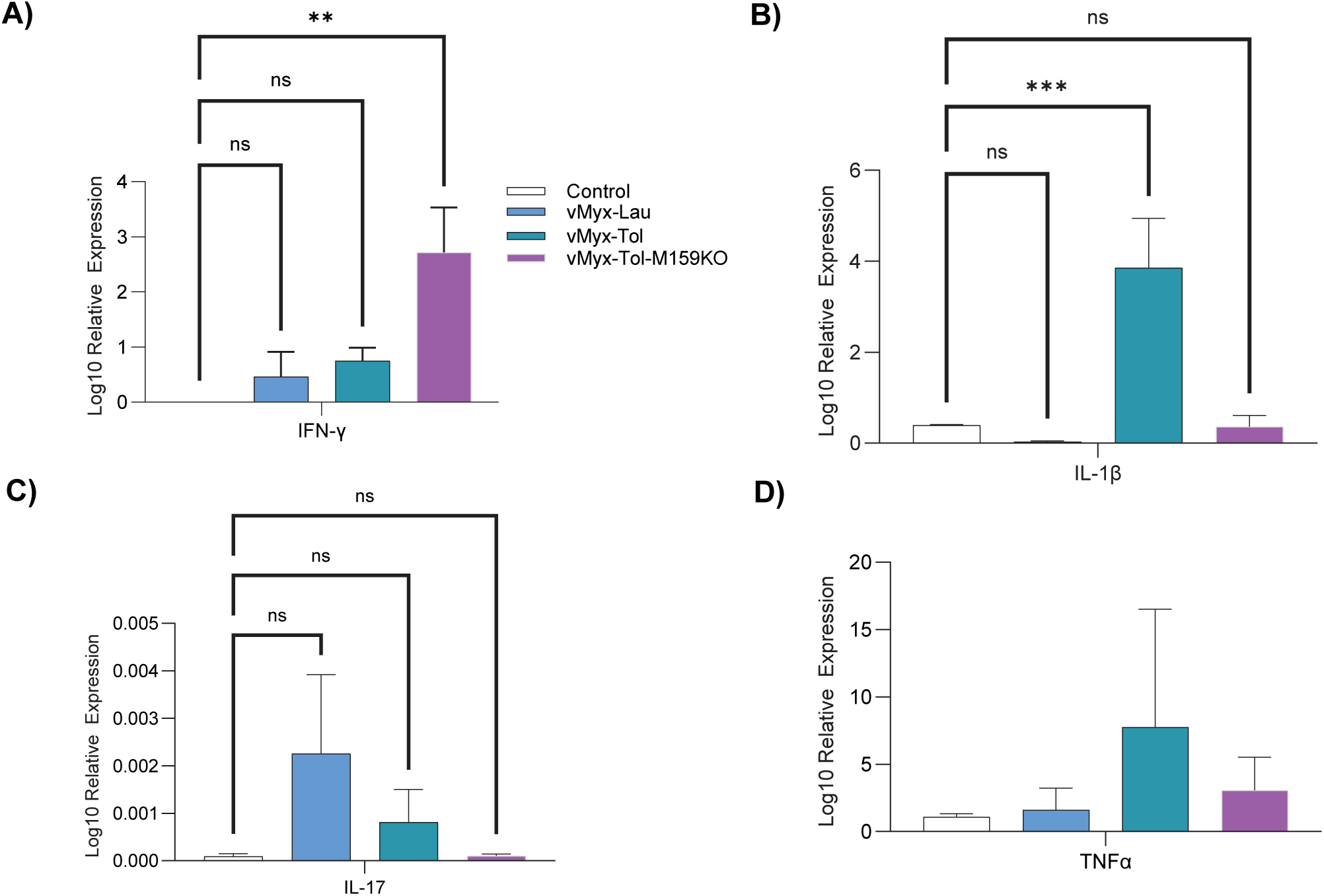
Analysis of gene expression (RT-qPCR) in primary skin lesions. Changes in A) IFNɣ, B) IL-1β, C) IL17, and D) TNFα expression were presented as a ratio of target gene vs. reference gene (GAPDH) relative to the expression in control samples.

